# Theoretical limits on the speed of learning inverse models explain the rate of adaptation in arm reaching tasks

**DOI:** 10.1101/2022.09.07.506913

**Authors:** Victor R. Barradas, Yasuharu Koike, Nicolas Schweighofer

## Abstract

An essential aspect of human motor learning is the formation of inverse models, which map desired actions to motor commands. Inverse models can be learned by adjusting parameters in neural circuits to minimize errors in the performance of motor tasks through gradient descent. However, the theory of gradient descent establishes an upper limit on the learning speed, above which learning becomes unstable. Specifically, the eigenvalues of the Hessian of the error surface around a minimum determine the maximum speed of learning in a given task. Here, we use this theoretical framework to analyze the speed of learning in different inverse model learning architectures in a set of isometric arm-reaching tasks. We show theoretically that, for these isometric tasks, the error surface and, thus the speed of learning, are determined by the shapes of 1) the force manipulability ellipsoid of the arm and 2) the distribution of targets in the task. In particular, rounder manipulability ellipsoids generate a rounder error surface, allowing for faster learning of the inverse model. Rounder target distributions have a similar effect. We tested these predictions experimentally in a virtual quasi-isometric reaching task with a visuomotor transformation. The experimental results were consistent with our theoretical predictions. Furthermore, our analysis accounts for the speed of learning in previous experiments with incompatible and compatible virtual surgery tasks, and in visuomotor rotation tasks with different numbers of targets. By identifying aspects of a task that influence the speed of learning, our results provide theoretical principles for the design of motor tasks that allow for faster learning.

**Author Summary:** When facing a new or changing environment, humans need to learn a new internal inverse model to generate fast and accurate movements. Different computational architectures based on supervised learning via gradient descent have been proposed to explain the acquisition of these inverse models. Although the theory of gradient descent and its results regarding the speed of learning are well developed in Machine Learning, this framework has seldom been applied to computational models of human motor learning. In this study, we found that applying this theoretical framework to a set of isometric reaching tasks clearly reveals aspects of the motor task that can speed up or slow down learning. In particular, we found that, in isometric tasks, the force manipulability ellipsoid of the arm and the distribution of force targets determine the speed of learning. These theoretical results allowed us to generate testable hypotheses about the speed of learning in different motor task conditions, which we successfully confirmed experimentally. We believe that our methods and results could open new lines of research to systematically identify aspects of motor learning tasks that can be exploited to enhance the speed of learning and to design new tasks that are easy to learn.

## Introduction

Humans are capable of producing fast and accurate movements to skillfully execute a variety of motor tasks. To accomplish such tasks, the central nervous system (CNS) uses feedforward mechanisms, as it cannot rely solely on delayed sensory feedback to guide the execution of movements [1,2]. Growing evidence indicates that the CNS generates feedforward motor commands via inverse models, which compute motor commands to achieve a desired sensory consequence given the current state of the body and environment, [3–8].

Inverse models must be able to adapt to changes in the environment or the body to maintain successful task execution. This adaptation faces two main challenges: 1) errors in task space do not directly inform how motor commands should be adjusted to eliminate the errors, and 2) the models are generally not unique because of the redundancy of the motor system (more motor commands than task variables). Accordingly, several computational models such as direct inverse modeling [9,10], distal learning [11], and feedback error learning [12] have been proposed as mechanisms to solve these challenges. These models differ in their architectures and mechanisms to adequately relate task errors to changes in motor commands. However, despite differences in their theoretical underpinnings, these models are all learned based on attempts to minimize an error quantity, such as performance error (distal learning) or motor error (direct inverse modeling and feedback error learning). Thus, from a computational perspective, the inverse learning problem can be formulated as a function fitting problem (mapping from desired motor outcomes to motor commands), where the strengths of neural connections in the inverse model circuit are the parameters that are tuned to fit the function.

It is now well accepted that the CNS encodes movement errors in the cerebellum, suggesting that the errors are used to drive motor adaptation in a supervised learning fashion [7,13–19]. This implies that the CNS uses some form of gradient descent on the errors to tune the parameters in the inverse model. In gradient descent, the parameters are adjusted in small steps in the direction which minimizes the error locally, given by first order derivatives of the error function with respect to the parameters. Thus, learning can be visualized as a path of successive steps in the multi-dimensional space of inverse model parameters. This path lies on a surface composed by the values of the error at each point in the inverse model parameter space.

Theoretical results in statistical learning establish that the shape of the error surface influences the speed of learning [20]. Round or isotropic surfaces allow faster learning because the gradient points in the general direction of the minimum of the error function, resulting in a relatively straight path to the minimum (see also [21,22]). In contrast, less round or anisotropic surfaces bring about slower learning because in large regions of the parameter space the gradient is misaligned with the direction of the minimum. The shape of the error surface can be described locally by the eigenvalues of a Hessian matrix, which contains the second order derivatives of the error with respect to the inverse model parameters [20].

Here, we provide a theoretical analysis of the speed of learning an inverse model using gradient-following optimization by analyzing the relationship between the Hessian matrix of the learning system, the goals of the task and the forward physics of the musculoskeletal system. In particular, we analyze learning in isometric arm tasks with visuomotor transformations, which change the relationship between produced forces and visual feedback, and musculoskeletal transformations, which change the role of muscles in force production. In these simple motor learning tasks, the CNS can be conceptualized as an inverse model represented as a feedforward neural network. In addition, the visuomotor and musculoskeletal transformations and the musculoskeletal system constitute the forward physics of the arm, which can be modeled altogether as a linear mapping [23,24]. As a result, the system composed of the inverse model and the forward physics of the arm (the musculoskeletal system and a transformation) can be viewed mathematically as a single feedforward network. We demonstrate that the Hessian matrix for this composite network can be readily computed, allowing us to make predictions about the speed of learning an inverse model for these tasks.

Analyzing the Hessian of the composite network allows us to identify how the goals and forward physics of the task influence the speed of learning under different transformations. In particular, we show that both 1) the shape of the distribution of target forces, and 2) changes in the force manipulability ellipse of the arm (the ability to generate end-effector forces given a joint configuration [25]) brought about by different transformations can limit the speed of learning.

In addition, we show that our theoretical results are valid for a range of assumptions about learning inverse models. Specifically, our results generalize to different learning architectures (distal learning and direct inverse learning), to different inverse model architectures (radial basis function networks and multi-layer perceptron networks), and to different gradient-following algorithms (backpropagation and node perturbation [26,27]). Importantly, we also show that our results are compatible with the theory of optimal control in motor learning, where the objective of learning the minimization of both errors and effort [28,29].

We then confirm our theoretical predictions via two experiments that tested the role of the force manipulability ellipse and the target distribution on the speed of learning a quasi-isometric task in a virtual environment. In the first experiment, we compare the speed of learning a visuomotor rotation with isotropic or anisotropic manipulability ellipses. In the second experiment, we compare the speed of learning a visuomotor rotation with targets in a circular or elliptical configuration.

Finally, we use our theoretical framework to account for differences in the speed of learning in two previous studies. First, we study a virtual surgery task, a type of musculoskeletal transformation in which the pulling forces of muscles in a mapping between muscle activations and cursor positions are changed, simulating tendon transfer surgeries [23]. Our results suggest that the shapes of the manipulability ellipses of the system after applying the virtual surgeries account, at least in part, for the differences in learning speed. Second, we address the speed of learning in visuomotor rotation experiments with different number of targets [30]. We show that our framework accounts for these differences in learning speed because the number of targets has a direct influence on the shape of the distribution of targets.

## Methods

### Computational modeling framework

#### Overview

We propose a computational framework to study the speed of motor learning during isometric arm reaching tasks, including visuomotor transformations and compatible and incompatible virtual surgeries [23]. We focus exclusively on feedforward control, which enables us to model the tasks as a static mapping between target and output forces, and to ignore transient forces. In the main text, we derive theoretical results based on the distal learning architecture [11] (Figure 1a), in which a forward model of the arm is used to estimate the relationship between task errors and motor commands to learn the inverse model. We then show in Appendix 1 that our framework generalizes to direct inverse modeling [9,10]. Note that feedback error learning is not applicable to this feedforward control isometric task.

**Figure 1.**
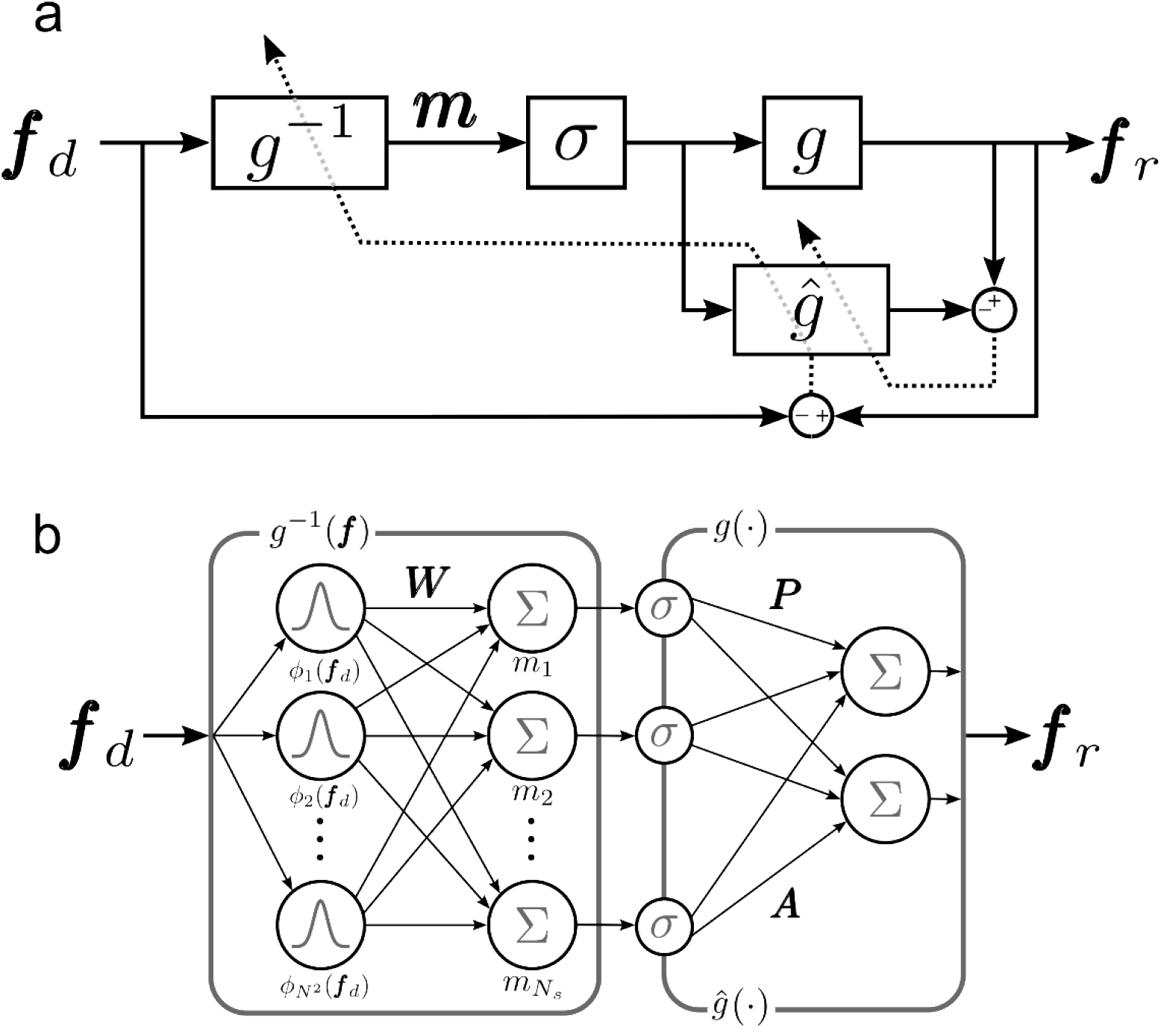
Structure of computational model for the distal learning of isometric arm tasks in which the inverse model is implemented with an RBF network **a**. Model diagram. The CNS learns an inverse model *g*^-1^ of the motor system *g* by using a forward model *ĝ* to update *g*^-1^ based on the minimization of the error ***f***_d_ – ***f***_r_. **b**. Structure of the inverse model *g*^-1^ and the motor system *g*. The input to *g*^-1^ is an RBF representation of a 2D force. The inverse model is a single-layer network and is fully defined by ***W***, the weights of the connections between the input and the output units. The output of *g*^-1^ is the motor command ***m***, which is the input to *g*. The motor system *g* and the forward model *ĝ* are linear transformations ***P*** and ***A***, respectively, of ***m*** after a non-linear transformation *σ*. Note that the CNS and motor system as a whole can therefore be seen as a two-layer network with linear outputs.

For simplicity, we use a model of the upper extremity that simulates production of isometric planar forces to reach different targets in a virtual environment. Because the tasks are isometric, the biomechanics of the arm can be approximated adequately as a linear mapping between motor commands and output forces [23,24,31,32].

#### Model structure and implementation

The model learns an inverse model *g*^-1^ of *g*, the motor system after applying a transformation *T*, which represents a visuomotor transformation or a virtual surgery. The inverse model *g*^-1^ receives a target 2-D force ***f***_d_ to generate a motor command ***m***, which can represent joint torques, muscle activations or muscle synergy activations [32]. Then, the motor command ***m*** propagates through the forward physics of the motor system *g*, producing a realized force ***f***_r_ (Figure 1a). Note that, despite its non-biological plausibility, we also consider the case in which ***m*** represents joint torques because it allows a totally analytical derivation of the properties of the speed of learning in the system, which provides insight to the more complex cases where ***m*** represents muscle or synergy activations. Specifically, the motor system *g* is represented as:

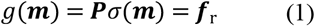

where ***P*** represents the forward physics of the isometric task, and ***m*** is an *N*_m_-dimensional motor command vector produced by the inverse model (joint torques, muscle activations or muscle synergy activations). Because the task is isometric, forces can be modeled as a linear function of the motor command ***m*** (or the transformed motor command *σ*(***m***)) [23,24,31,32]. Thus, if the motor system is unperturbed, then ***P*** = ***M***, where ***M*** is a matrix containing a linear mapping between motor commands and planar forces (2 × *N*_m_) [23,32]. A transformation *T*, such as a visuomotor transformation or virtual surgery, can alter the forward physics of the task such that ***P*** = *T*(***M***) (see Motor command to force mapping, Visuomotor transformations, and Virtual surgeries subsections below for details). If ***m*** represents joint torques the activation function *σ*(***m***) is a linear function. Otherwise, it is a non-linear function that maps each element of ***m*** into a non-negative muscle or synergy activation. The non-linear function can be a sigmoid or a rectified linear unit (RELU) function.

We first consider that the inverse model *g*^-1^ is learned via a radial basis function (RBF) network, as they are biologically plausible [33], and simplify theoretical derivations. With this inverse model architecture, and for isometric tasks, the structures of *g*^-1^ and *g* and the connections between them allow us to view *g*^-1^ and *g* as a single neural network with one hidden layer and linear outputs (Figure 1b). We note however that our approach is not limited to the RBF architecture, and show that it extends to multi-layer perceptron networks, for instance (see *Generalizability of the results to muscular models, realistic error function, and neural network structure* section below). In the RBF network, the inverse model is given by:

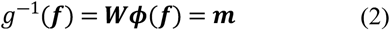

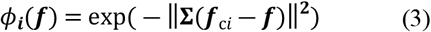

where ***W*** is a matrix of weights (*N*_m_ × *N*^2^), ***ϕ***(***f***) is a vector of *N*^2^ two-dimensional RBFs *ϕ_i_*(***f***) evaluated at ***f***, ***f***_c*i*_ are the centers of the RBFs placed on a *N* × *N* square grid centered on the origin of the force space, and **Σ** determines the shape of the RBFs.

The distal learning model includes an internal forward model *ĝ*, which is used to estimate the relationship between task errors and changes in the motor commands produced by *g*^-1^ [11]. The forward model *ĝ* uses an efferent copy of ***m*** to produce 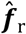, an estimate of ***f***_r_, to compute a sensory prediction error 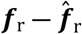. Since the forward physics of the arm are linear, we define *ĝ* as a linear operator on *σ*(***m***):

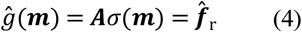

where ***A*** has the same dimensions as ***P***, so that *ĝ* has the same structure as *g* (eq. 1).

#### Learning of the inverse model

Learning in the isometric tasks aims to minimize the task performance error given by the cost function

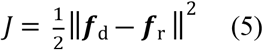

In distal learning, the learner uses the forward model *ĝ* to compute an approximate gradient of *J* because exact knowledge of *g* is not available. Assuming that *ĝ* is a good enough representation of *g*, learning converges to the minimum of *J* [11]. As stated above, we can view *g*^-1^ and *ĝ* as a composite neural network (Figure 1b). Therefore, the approximate gradient for the network can be computed using methods such as backpropagation, stochastic methods like weight or node perturbation [26,34], or other reinforcement learning-based methods like REINFORCE [27]. Here, we used backpropagation to illustrate the properties of the speed of learning under gradient-following algorithms. However, we also show that our results do not depend on this choice, as they are also valid for the node perturbation algorithm (Appendix 2).

The learning procedure starts with the forward propagation of a given ***f***_d_ through *g*^-1^ and the actual motor system *g*, resulting in ***f***_r_ and a task performance error ***f***_d_ – ***f***_r_. Then, the task performance error is backpropagated through *ĝ* and *g*^-1^, which provides an approximation of ∇_***W***_*J*, the gradient of *J* with respect to the weights ***W*** of *g*^-1^ (eq. 2). Lastly, ∇_***W***_*J* is used to update ***W***:

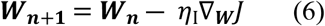

where ***W_n_*** is the weight matrix at the current trial, ***W*_*n*+1_** is the updated weight matrix, and *η*_I_ is the learning rate parameter.

We assume that the forward model is learned at a much faster rate than the inverse model, and that learning of the forward model is perfect. That is, we set ***A*** = ***P*** = *T*(***M***) according to the transformation T used in the task. This is justified because learning forward models is much simpler than learning inverse models, and inverse models can be learned via distal learning even when using imperfect forward models [11]. This assumption allows us to apply the methods described in the next section to estimate the limits on the speed of learning the inverse model *g*^-1^ when the forward physics are subject to *T*.

#### Estimation of upper limit on learning speed

In this section we demonstrate that the speed of learning in the isometric task is limited by the shapes of the force manipulability ellipse and the distribution of targets in the task.

A given cost function can be approximated locally around the minimum of the cost function as a second order Taylor expansion [20]:

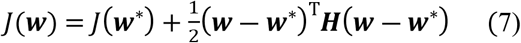

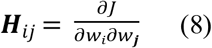

where ***w*** is a vector of weights, ***w***\* is the vector of weights at the minimum of the cost function *J*, and ***H*** is the Hessian of *J*, a matrix of second order derivatives of *J* with respect to ***w***. In this second order approximation, the gradient of *J* is ∇_***W***_*J* = ***H***(***w*** – ***w***\*). Therefore, the update equation of ***w*** using gradient descent can be approximated by the following learning rule

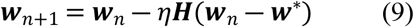

where *η* is the learning rate parameter. Previous work has demonstrated that in the vicinity of ***w***\*, there is an upper limit on the value of *η*, beyond which updates to ***w*** become unstable. This value of *η* also determines the slowest time constant of learning, which limits the speed of convergence to ***w***\* and is a function of *λ*_***H***min_/*λ*_***H***max_, the ratio of the largest and smallest eigenvalues of ***H*** [20] (See Appendix 3 for a derivation of this result). From a geometric point of view, ***H*** describes the shape of the cost function *J* around ***w***\*. Namely, it can be shown that the contours of constant *J* are ellipsoids, and that the sizes of the axes of these ellipsoids are proportional to the square roots of the non-zero eigenvalues of ***H***. Therefore, if both eigenvalues *λ*_***H***min_ and *λ*_***H***max_ have similar magnitudes, the shape of the cost function *J* is relatively round, allowing faster convergence to the minimum. On the other hand, a large disparity in the magnitudes of *λ*_***H***min_ and *λ*_***H***max_ corresponds to a highly anisotropic *J*, producing slower convergence to the minimum (see Appendix 3 for an illustration of this effect).

Based on these theoretical results relating the speed of learning and the Hessian ***H*** of the learning system, we aimed to derive ***H*** and its eigenvalues in the distal learning system for isometric tasks. In particular, we investigate how the visuomotor and musculoskeletal transformations affect the structure of ***H*** and its eigenvalues, allowing us to estimate the limits on the speed of learning for each task. For mathematical simplicity, let us first consider the case in which the activation function *σ*(***m***) is linear, that is *σ*(***m***) = ***m***. This can be interpreted as ***m*** representing a joint torque vector, and ***P*** representing a mapping between joint torques and end-point forces of the limb (i.e. the inverse Jacobian of the limb). As indicated in the *Learning of the inverse model* section above, we assume that *ĝ* learns a perfect model of the perturbation as soon as the perturbation is introduced, namely, *ĝ* becomes equal to *g*. This makes the distal learning procedure equivalent to a classic linear supervised learning problem in which the gradient can be obtained analytically. This allows us to compute the Hessian ***H*** analytically, and to estimate the limits on the speed of learning in terms of the goals and forward physics of the tasks.

Using eq. 2 and 5, the cost function is:

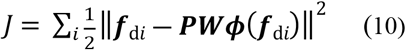

where ***f***_d*i*_ corresponds to each target in the task, with *i* = 1, …, *N*_T_, and *N*_T_ is the number of targets.

Using standard matrix algebra (see Appendix 4), the gradient ∇_***W**J*_ is

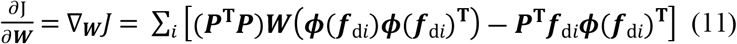

and the Hessian ***H_W_*** is

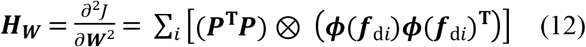

where the symbol ⊗ denotes the Kronecker product. The derivation of eq. 12 is provided in Appendix 4. We now use the associative property of the Kronecker product:

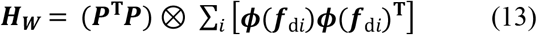

The term on the right side of the Kronecker product is a scaled sample covariance matrix of the RBF representation of targets ***≠***(***f***_d*i*_):

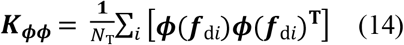

where *N*_T_ is the number of targets in the task. Therefore:

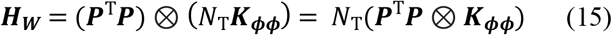

The eigenvalues of the Kronecker product of two matrices are the set of products between each eigenvalue of the first matrix and each eigenvalue of the second matrix [35]. That is, 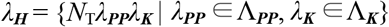 where Λ_***PP***_ and Λ_***K***_ are the sets of eigenvalues of ***P***^T^***P*** and ***K_ϕϕ_***, respectively. Because both ***P***^T^***P*** and ***K_ϕϕ_*** are positive semi-definite (all eigenvalues are non-negative) the largest and smallest eigenvalues of ***H*** are *λ*_***H***max_ = *N*_T_*λ*_***PP***max_*λ*_***K***max_ and *λ*_***H***min_ = *N*_T_*λ*_***PP***min_*λ*_***K***min_, respectively. Thus,

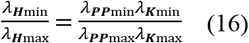

Thus, the ratio *λ*_min_/*λ*_max_ for ***H_W_*** is a function of the *λ*_min_/*λ*_max_ ratios of ***P***^T^***P*** and ***K_ϕϕ_***. The matrix ***P***^T^***P*** is related to the properties of the motor system *g*, and how it interacts with the perturbation *T*. The matrix ***K_ϕϕ_*** is related to the distribution of target forces ***f***_d_.

Next, we show how the ratio *λ*_***PP***min_/*λ*_***PP***max_ is directly related to the force manipulability ellipse of the arm. The force manipulability ellipse is a mapping from a unit ball of joint torques to endpoint forces. It characterizes the force generation capabilities of the arm, showing that the arm can more readily produce forces along the major axis of the ellipse than in other directions [25]. If ***P*** corresponds to a mapping between joint torques and end-point forces, the mapping can be described by the force manipulability ellipse. The axes of the force manipulability ellipse have sizes that correspond to the singular values *s*_***P***_ of ***P*** [25]. It is also known that 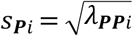. Thus,

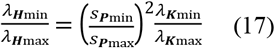

The force manipulability ellipse concept can be generalized to the case where ***P*** corresponds to a mapping between muscle or synergy activations and forces. In this case the ellipse is the image of a unit ball of muscle or synergy activations in the space of end-point forces.

We have therefore demonstrated analytically that 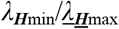 can be controlled by independently adjusting the force manipulability ellipse of ***P*** (varying the ratio of the magnitudes of the ellipse axes), or by adjusting the distribution of targets ***f***_d_ or other RBF parameters in the ***ϕ*** function. Notice that the transformation *T* in ***P*** can alter the shape of the force manipulability ellipse. This happens if *T* is a scaling transformation or a virtual surgery, as described in the *Simulation and Experimental methods* sections below.

#### Generalizability of results to realistic error function, muscular models, and neural network structure

The theoretical results derived above were obtained for a learning system that minimizes performance errors. However, there is ample evidence that suggests that the CNS actually minimizes a trade-off between performance errors and effort during learning [28,29,36]. In Appendix 5, we show that our theoretical results generalize to learning systems that optimize a trade-off between the performance error and the effort term, which penalizes large motor commands.

Additionally, the above results are based on the idealized case of a linear system with joint torques as the motor command. In Appendix 6 we demonstrate that introducing the nonlinearity *σ*(***m***) to the motor command produces qualitatively similar results to the linear case. That is, we show that our theoretical results are also useful to model the speed of learning in systems with pulling muscles.

Finally, although we defined above the inverse model *g*^-1^ as an RBF network, the above results can generalize to other inverse model architectures. For instance, the RBF network in the inverse model can be replaced by a typical multi-layer feedforward network or any other function approximator. To see why this is the case, we can visualize the new inverse model to include a function *h* on the network input with tunable parameters ***w***_pre_ such that

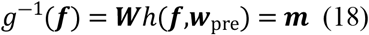

This functional form also encompasses the inverse model in eq. 2, as *h*(***f,w***_pre_) can describe the output of the RBF units. Therefore, if we assume that a set of valid parameters ***w***_pre_ is known prior to learning and learning only occurs on ***W***, then all the above arguments about the speed of learning are also valid for the new architecture. This idea is similar to our treatment of the RBF network, where the RBF unit parameters (center and shape) are not considered for the calculation of the gradient. Of course, the parameters ***w***_pre_ are generally not known a priori, so they must be learned using gradient descent or other methods. However, it follows that the dynamics of learning ***W*** must be influenced by the shapes of the force manipulability ellipse and the target distribution in the same way that we have described above.

### Simulation methods

Based on the theoretical results of our computational framework, we simulated learning in four isometric arm tasks: 1) anisotropic scaling task, 2) anisotropic target distribution task, 3) virtual surgery task, and 4) target number task. In the anisotropic scaling and virtual surgery tasks we control the shape of the manipulability ellipse of the arm in different conditions. In the anisotropic target distribution and target number tasks, we control the target distributions in different conditions. The virtual surgery and target number tasks were included to explain experimental findings reported in previous studies in the context of our theoretical results. We simulated all tasks using the distal learning and direct inverse modeling paradigms (Appendix 1). In the previous section we described a network in which the motor commands are joint torques for illustration purposes. However, our main interest is using a musculoskeletal system for a more accurate representation of the CNS. Therefore, we used sigmoidal activation functions in the outputs of the inverse model so that the motor command corresponds to muscle activations (Appendix 6). We also considered network architectures in which muscle synergies are explicitly included in the model (Appendix 7). This is important, because previous results in the virtual surgery task attribute differences in the speed of learning between conditions to the presence of muscle synergies in the CNS [23]. Finally, we also considered a cost function that minimizes a trade-off between error and effort in the task in all simulations (Appendix 5, 6).

We ran all four simulated tasks with 15 simulated participants. Each simulated subject consists of a mapping between muscle activations and planar forces (***M***) and a set of muscle synergies (***S***). We used experimental data from 15 human participants during an isometric reaching task to obtain ***M*** and ***S*** for each simulated participant as described in Barradas et al. 2020. This allowed us to create simulated participants that produce realistic muscle activations (Appendix 8).

We also computed 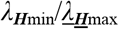, the ratio of the smallest and largest eigenvalue of ***H***, for each simulated subject in each task according to the model structure described above using eq. A6.11 and as described in Appendix 6.

#### Motor command to force mapping

Each simulated participant was defined by a mapping between motor commands ***m*** and forces ***f***_r_. This mapping was obtained from a real human participant in a previous study [32], as described in Appendix 8. We focus on two cases: ***m*** represents activations of muscles that cross the shoulder and elbow, or ***m*** represents muscle synergy activations of the same muscles. In both cases, the mapping is defined by ***P*** in eq. 1. In the case of a muscle activation to force mapping ***P*** = ***M***, where each column of ***M*** indicates the force produced by each muscle, which is scaled by the elements of *σ*(***m***). Notice that the nonlinearity *σ* ensures that ***m*** will produce only pulling forces as defined by ***M***. The ***M*** mappings taken from the real human participants had 10 muscles (*N*m = 10). We describe the simulation methods related to synergy activations in Appendix 7.

#### Anisotropic scaling task

Based on the theoretical results in eq. 17, we designed a task with visuomotor transformations that control the shape of the force manipulability ellipse of ***P*** to show its effects on the speed of learning. In our models, visuomotor transformations modify the motor command-force mapping ***P*** by applying a linear transformation in force space:

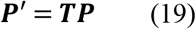

where ***T*** is a 2 × 2 matrix that constitutes the visuomotor transformation. We designed ***T*** as the combination of a rotation and a scaling transformation. We use a rotation in addition to the scaling so that we can observe learning even in the absence of scaling:

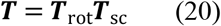

where ***T***_rot_ is a 30° counterclockwise rotation, and ***T***_sc_ is a linear transformation that alters the shape of the manipulability ellipse, given by the ratio of the singular values *s*_min_/*s*_max_ of ***P*’**. The ***T***_sc_ transformation is a composition of a rotation and a scaling that allows to directly control *s*_min_/*s*_max_. Because the axes of the manipulability ellipse of **P** are not necessarily aligned with the horizontal and vertical directions, ***T***_sc_ must first align ***P*** to the axes of the ellipse:

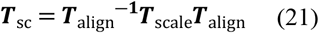

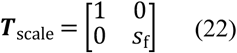

Therefore, the ***T***_sc_ transformation first rotates ***P*** according to a rotation ***T***_align_ such that one of the left singular vectors of ***P*** corresponding to *s*_max_ becomes aligned with the x axis, then scales the aligned ***P*** according to a scaling ***T*** scale such that *s*_min_/*s*_max_ changes, and finally, rotates ***P*** again according to the inverse rotation ***T***_align_^-1^ such that the left singular vector of ***P*** returns to its original orientation. The transformation ***T***_scale_ is defined by *s*_f_, a scaling factor. We created three conditions in which *s*_f_ = [0.2 0.5 1].

#### Anisotropic target distribution task

Based on the theoretical results in eq. 17, we designed a task with targets defined according to distributions of different shapes to show their effects on the speed of learning. The task was a 30° counterclockwise visuomotor rotation. We defined two different target distributions. In the first one, targets were placed uniformly on the outline of a circle on the horizontal plane, constituting an isotropic distribution. In the second one, targets were placed on the outline of an ellipse, constituting an anisotropic distribution. The targets in the anisotropic distribution were defined by applying a scaling transformation on the targets of the isotropic distribution:

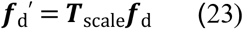

where ***f***_d_ are the target forces in the isotropic distribution, ***f***_d_’ are the target forces in the anisotropic distribution, and ***T***_scale_ is a scaling transformation as defined in eq. 22. The value of the scaling factor *s*_f_ was set to 0.2.

#### Virtual surgery task

Following the methods described in Berger et al. 2013, we designed virtual surgeries that were either incompatible or compatible with the muscle synergies ***S***_ext_ extracted from the models (see Appendix 7). A virtual surgery modifies the forward physics of the arm (***P***) by applying a linear transformation in muscle space:

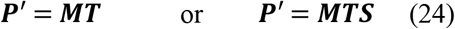

where ***T*** is an orthonormal *N*_m_ × *N*_m_ matrix that constitutes the transformation or virtual surgery, and ***S*** is a *N*_m_ × *N*_s_ matrix representing the synergies embedded in the model. *N*_s_ is the number of synergies in the model. The first equation corresponds to the case in which ***m*** are muscle activations, and the second equation to the case in which ***m*** are muscle synergy activations.

Incompatible virtual surgeries are designed such that muscle activations produced by ***S*** are restricted to generate forces along only one dimension of the force space, while the resulting muscle activation-force mapping ***MT*** spans the whole force space. Therefore, theoretically, any force can be produced by a new combination of muscle activations, but in practice, forces are biased towards one dimension of the plane. In contrast, compatible virtual surgeries are designed such that muscle activations produced by ***S*** span the whole force space. We show that incompatible surgeries produce highly anisotropic manipulability ellipses of ***P***, whereas compatible surgeries do not affect the shape of the manipulability ellipse. Therefore, both kinds of surgeries have different effects on the speed of learning in the task.

For each simulated subject, we designed an incompatible and a compatible surgery based on the simulated subject’s ***M*** and extracted ***S***_ext_. We designed both surgeries according to the methods described in [23]. The incompatible surgeries were built by finding ***T***_I_ that maps muscle activations in the column space of ***S***_ext_ into the null space of ***M***. In contrast, the compatible virtual surgeries were built by finding ***T***_C_ that transforms the muscle activations into different muscle activations outside of the null space of ***M***. Details can be found in [23]. We randomly generated incompatible and compatible surgeries and applied them to the forward physics of the system to calculate the initial average error in force direction under each surgery. We used eight radially and uniformly distributed force targets. We selected surgeries that produced an initial average error in force direction close to 60°. This was decided based on previously reported results for the mean initial error during incompatible and compatible virtual surgeries [23]. We followed the experiment schedule defined in [23].

#### Target number task

We simulated a visuomotor rotation reaching task, in which target sets with different numbers of targets are presented. This simulated task is based on a previous experiment [30]. We show that different numbers of targets produce differences in the eigenvalues of the covariance matrix of targets ***K_ϕϕ_***, which influences the speed of learning. We used 3 target sets with 1, 4 and 8 targets each. The targets in the 4- and 8-target sets were radially and uniformly distributed. The visuomotor rotation was a counterclockwise 30° rotation. We followed the experiment schedule defined in [30].

#### Simulation procedure

We ran simulations of each experiment for 15 simulated participants based on mappings between muscle activations and planar forces ***M*** and muscle synergy matrices ***S*** computed in a previous study [32]. We trained each simulated subject to adequately perform the baseline task without perturbations while producing plausible muscle activations (see Appendix 8 for details). Each simulation trial was defined as one cycle of target presentation, force production, and a learning step. The simulations followed the protocol defined in the corresponding experiment (number of baseline and perturbation trials defined in the *Experimental methods* section, in [23] and in [30]), except for counter-perturbation and washout phases, which we did not simulate.

The model takes the learning rate of the inverse model *η*_I_ as a parameter (eq. 6). We defined a set of learning rate values (*η*_I_ = [0.0001, 0.00015, 0.0002, 0.0005, 0.00075, 0.001]) and ran simulations for each value. For each value of η_I_ and simulated subject, we ran 10 different simulations with a different target order. The target order in each simulation was pseudo-randomly generated within cycles of the defined targets. We averaged the learning curves corresponding to the error in force direction obtained from each set of these 10 simulations. We then further averaged the error in direction for every sequence of eight targets to obtain smooth learning curves (except for the multi-target experiment, in which there are only 18 trials per condition). For each simulated subject, we selected the value of *η*_I_ that best fit the mean experimental results in the corresponding task, as measured by the root mean square error (RMSE) between the subject’s and the mean experimental learning curves. Note that for a given simulated participant, the value of *η*_I_ was the same for all simulated tasks. Finally, we averaged the resulting learning curves across all 15 simulated participants.

### Experimental Methods

Based on the insights provided by our computational framework, we hypothesized that anisotropy in the force manipulability ellipse and in the distribution of targets would slow down motor learning with respect to more isotropic force manipulability ellipses and target distributions. To test these hypotheses, we performed experiments based on the anisotropic scaling and anisotropic target distribution tasks described in the Simulation methods section above. Experiments for the virtual surgery task and the number of targets task have been performed in previous studies [23,30].

#### Participants

Twenty-three right-handed participants [mean age, 27.6 yr (SD 5.7); 19 men] were included in the study after providing written informed consent. All 23 participants participated in Experiment 1, whereas a subset of 21 participants participated in Experiment 2. Additionally, we used data from 15 participants reported in a previous study [mean age, 27.9 yr (SD 8.8); 13 men] [32] to model learning in the isometric tasks. All procedures were approved by the Ethical Review Board of the Tokyo Institute of Technology.

#### Experimental setup

Because of the COVID-19 pandemic, which restricted in-person experiments, the experiments were implemented on an online platform (OnPoint) that participants accessed remotely using a laptop computer [37]. This online platform has been shown to produce similar results to in-person experiments in studies of adaptation to visuomotor transformations [37–39]. Participants were instructed to place the laptop on a table or desk and to sit comfortably in front of it with the laptop centered at the body midline (Figure 2a). A virtual environment was displayed on the laptop screen. The virtual environment consisted of a circular white cursor, a ring-shaped white target at the center of the screen, and several circular blue targets displayed on a black background.

**Figure 2.**
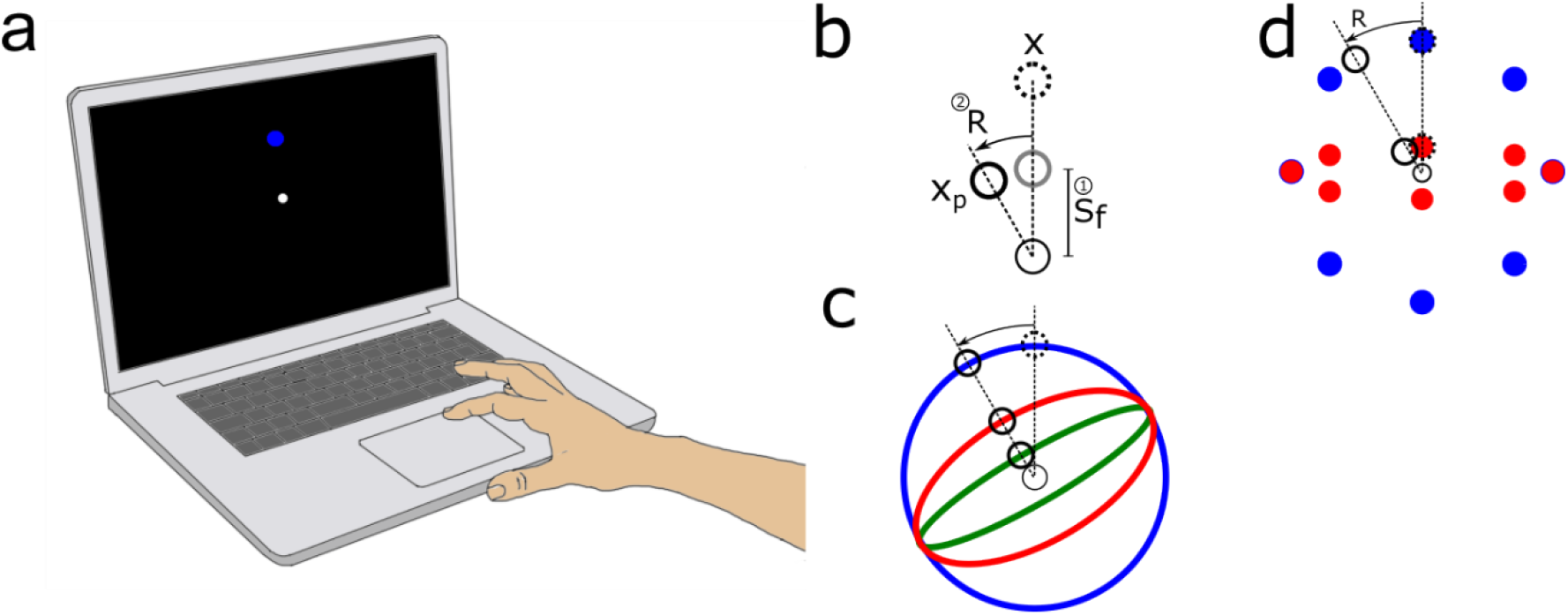
Experimental setup and protocols. **a**. Experimental setup. Participants performed a reaching task using the trackpad of a laptop computer. **b**. Experiment 1: anisotropic scaling. During the perturbation and counter-perturbation phases of experiment 1, visual feedback of the cursor was transformed by scaling the position of the cursor in the vertical axis of the screen by a factor s_f_ and rotating the resulting position 30° around the center of the screen in the counterclockwise or clockwise direction. **c**. Experimental conditions in experiment 1. Each participant performed the reaching task in three separate sessions. In each session the scaling factor *s*_f_ was different (1.0, blue; 0.5, red; 0.2, green) but the rotation angle was the same. **d**. Experiment 2: target distributions. Each participant performed the reaching task with a visuomotor rotation in two separate sessions. In each session the distribution of target positions was different. In the first condition targets were uniformly and radially distributed around the center of the screen. Thus, this condition is equivalent to the condition in experiment 1 in which *s*_f_ = 1 (blue). In the second condition the targets were arranged into an elliptical shape by scaling the vertical position of the targets in the first condition by *s*_f_ = 0.2 (red).

#### Experimental protocols

Participants used their right index finger and the built-in trackpad of a laptop computer to control the position of a cursor on the visual display. The experimental task required the displacement of the cursor from a center position to one of eight targets uniformly distributed around the center of the screen. Participants were instructed to make quick center-out movements with the cursor towards the target, and to attempt to stop inside the target. Participants were also informed that for some trials visual feedback of the cursor would be perturbed and not match their expected feedback. Participants were instructed to continue aiming toward the displayed target and to ignore the perturbation, which allows the observation of the implicit component of motor adaptation [40].

Each trial started by displaying the target at the central position, which roughly corresponded to the center of the trackpad. After the cursor was placed inside the central target for 500 ms, one of the targets appeared. Visual feedback of the cursor position was suppressed during the reaching movement and was only provided at the end of the trial as a static image of the cursor at the position where the cursor stopped. Visual feedback of the cursor at the end of the trial was held on the screen for 1 s, after which it was suppressed. Next, the central target and a white ring appeared. The radius of the white ring indicated the distance between the cursor and the central target and was used to eliminate direction-specific feedback of the cursor. Participants were asked to return to the central target by reducing the diameter of the ring. The central target turned into a white circle when the cursor was placed inside of it. After this, the next trial started.

#### Experiment 1: Anisotropic scaling

Participants completed three experimental sessions on different days, with 1 to 4 days between sessions, depending on the availability of each participant. In each session, participants performed a total of 43 target cycles (344 trials), divided into baseline, perturbation and counter-perturbation phases, which consisted of 5, 25 and 13 target cycles, respectively. Each target cycle contained a sequence of eight targets distributed uniformly around the center of the screen in a pseudorandom order. In the baseline phase, cursor feedback was veridical. In the perturbation phase, a composition of a 30° counterclockwise rotation and a scaling transformation was applied to the position of the cursor (Figure 2b). In the counter-perturbation phase, the rotation was switched to the clockwise direction, and the scaling transformation was the same as in the perturbation phase. The counter-perturbation was included to induce retrograde interference [41], minimizing possible savings in learning the rotation aspect of the perturbation in subsequent sessions. In a similar way to the simulated anisotropic scaling task, the perturbations were defined as:

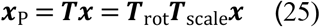

where ***x*** and ***x***_P_ are two-dimensional vectors containing the position of the cursor on the screen before and after applying the perturbation, ***T*** is a visuomotor transformation consisting of a scaling and rotation, ***T***_rot_ is a rotation matrix, and ***T***_scale_ is a scaling matrix as defined in eq. 22, The value of the scaling factor *s*_f_ in ***T***_scale_ was different in all three sessions (0.2, 0.5 or 1, Figure 2c), and the order of the sessions was counterbalanced across participants.

#### Experiment 2: Target distribution

A subset of the participants in Experiment 1 completed an additional experimental session after 2 – 4 weeks from the last session in Experiment 1. The trial structure in Experiment 2 was identical to that of Experiment 1, except for the position of the targets and the nature of the perturbation and the counter-perturbation. The positions of the targets were defined according to the anisotropic target distribution described in the *Simulation methods* section (scaling factor *s*_f_ = 0.2, Figure 2d). The perturbation was a 30° counterclockwise rotation of the cursor position, and the counter-perturbation was a 30° clockwise rotation, both with no scaling component.

The results of Experiment 2 were compared to the results of the condition in Experiment 1 in which *s*_f_ = 1. This is because this condition is equivalent to using a pure rotation and a round target distribution.

#### Learning rates

We measured the error in force direction between the target and the final cursor position as the outcome variable. Similar to the simulations, we averaged the error for all targets in a target cycle. For each subject and condition, we fitted the error curve with a first order exponential to obtain a learning rate parameter. We used the *fit* non-linear least squares fitting function in Matlab R2019b. We compared the computed learning rate parameter to 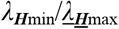 based on the Hessian of the network of the corresponding condition.

#### Statistical analysis

The main outcomes of the experiments were the degree of adaptation to the visuomotor transformations (initial error – final error in reaching direction) and the computed decay rate parameter of the exponential fit to the time course of the error in direction, estimated as indicated in the *Learning rates* section above. For Experiment 1, we tested the null hypothesis that the means of the outcome variables in the three anisotropic scaling conditions were equal by means of a repeated measures ANOVA. In cases where the outcome variables did not satisfy the normality assumptions of the ANOVA test, we performed a longitudinal non-parametric test using the nparLD package in R [42].We then performed post-hoc multiple comparisons using Bonferroni corrections. In Experiment 2, we tested the null hypothesis that the means of the outcome variables in the two target distribution scaling conditions were equal by means of a paired *t* test. In cases where the outcome variables did not satisfy the normality assumptions of the *t* test, we performed a Wilcoxon signed rank test. The significance threshold was set to *p* = 0.05. All analyses were performed in R 4.2.1.

## Results

### High anisotropy in manipulability ellipsoids is associated with lower degrees of adaptation to visuomotor transformations

Figure 3 shows the performance of a representative participant before, at the onset and at the end of the transformation block for all conditions in experiments 1 and 2. In experiment 1 (anisotropic scaling task), we found that scaling the manipulability ellipse had a statistically significant effect on the degree of adaptation to the visuomotor transformation (initial error – final error) [s = 1.0, initial error: 27.4° (SD 3.4), final error: 18.9° (SD 6.1); s = 0.5, initial error: 27.3° (SD 2.3), final error: 22.5° (SD 4.9); s = 0.2, initial error: 30.2° (SD 3.3), final error: 28.3° (SD 4.1); *F*(2,44) = 12.98, *p* < 1×10^-4^, repeated measures ANOVA]. Post hoc tests indicated that the degree of adaptation was significantly larger when no scaling factor was used (s = 1.0) [s = 1.0 vs s = 0.5, *p* = 0.027; s = 1.0 vs s = 0.2, *p* < 1×10^-3^]. However, there was no statistically significant difference in the degree of adaptation between scaling factors s = 0.5 and s = 0.2 (*p* = 0.12).

**Figure 3.**
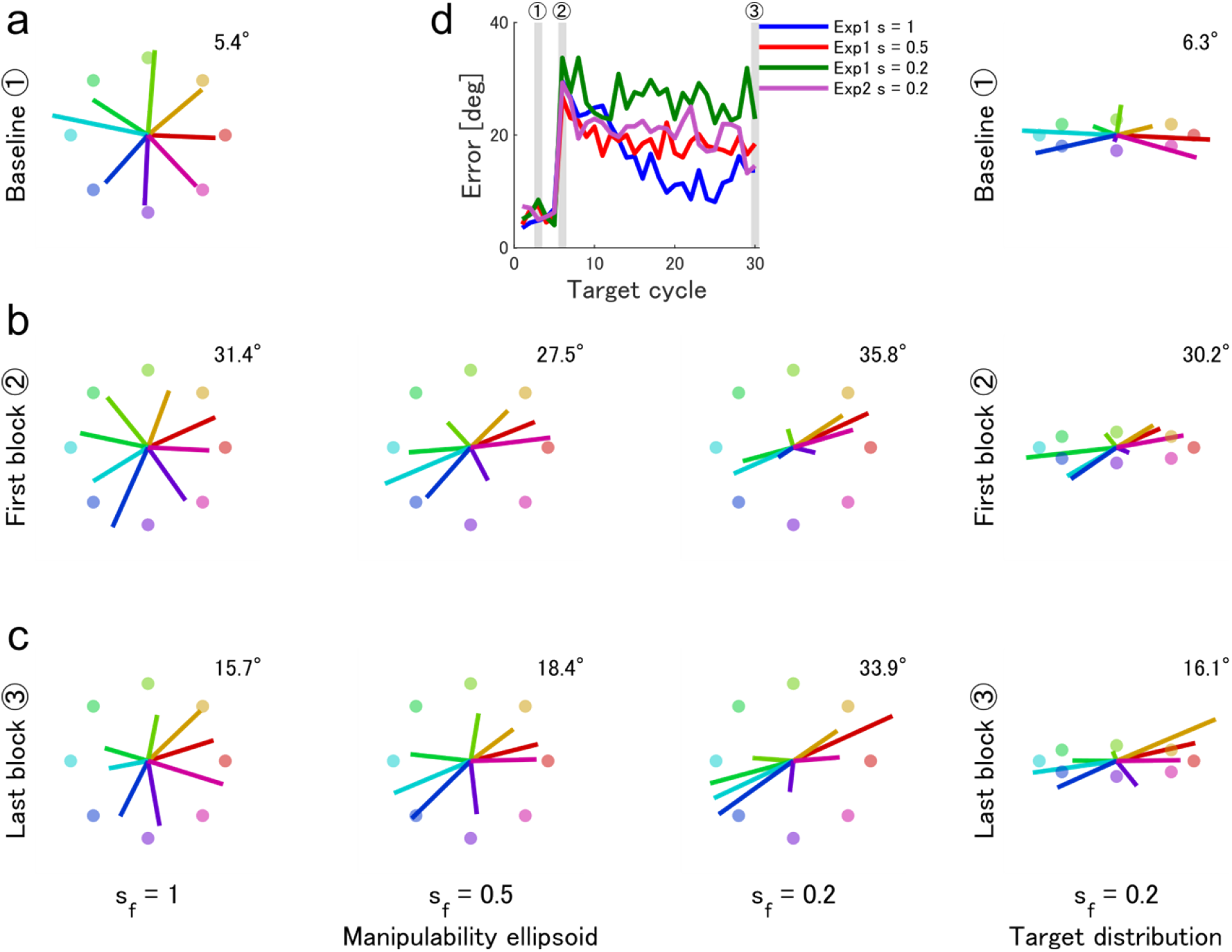
Cursor positions at the end of a target cycle before, at the onset, and at the end of training in all conditions of experiments 1 and 2 for a representative participant. **a**. Baseline block. **b**. First block of training. **c**. Last block of training. The mean absolute value of the error in final direction is indicated for each condition and target cycle. **d**. Performance error measured as the mean absolute value of the error in initial direction for one block, comprising 8 targets. Shaded areas indicate the blocks corresponding to the cursor positions in **a**, **b** and **c**.

Figure 4b shows the time course of learning as measured by the mean absolute error in the direction of the reach in all conditions of experiment 1. We found that scaling the manipulability ellipse had a statistically significant effect on the learning rate parameter of a single exponential fit to each participant’s learning curve (Figure 4a) [s = 1.0, learning rate: 0.0186 (SD 0.0196); s = 0.5, learning rate: 0.0077 (SD 0.0081); s = 0.2, learning rate: 0.0027 (SD 0.0036); *F* = 21.9, df = 1.77, *p* < 1×10^-5^, nparLD (data in condition s = 1.0 not normal according to Shapiro-Wilk test)]. Post hoc tests indicated that the learning rate parameter was significantly larger when no scaling factor was used (s = 1.0) [s = 1.0 vs s = 0.5, *p* < 10^-3^; s = 1.0 vs s = 0.2, *p* < 10^-5^]. Additionally, the learning rate parameter in scaling condition s = 0.5 was significantly larger than in condition s = 0.2 *(p* = 0.039). The differences in the extent of adaptation and learning rates in all the experimental conditions suggest faster learning rates for tasks with the least anisotropic virtual manipulability ellipsoids.

**Figure 4.**
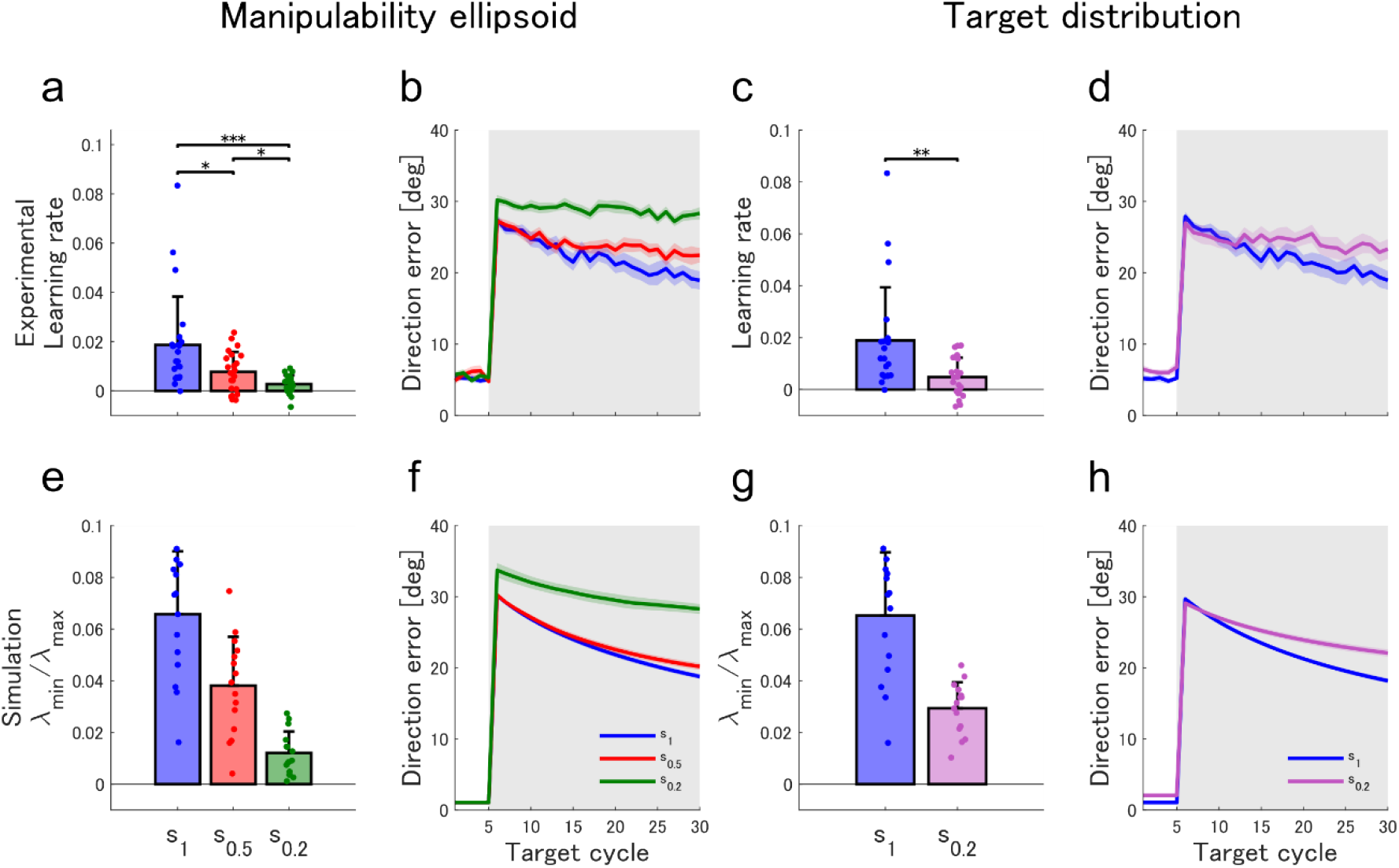
Experimental and simulation results in the anisotropic scaling and anisotropic target distribution tasks. **a**. Experimental learning rates in Experiment 1, and **c**. Experiment 2. We estimated the experimental learning rates by fitting the learning curves of each participant to simple exponentials. Bars indicate the mean, and error bars the standard deviation. Asterisks indicate significant differences between conditions: *** *p* < 0.001, ** *p* < 0.01, * *p* < 0.05. **b, d**. Experimental mean error in initial direction during learning in Experiments 1 and 2. **e**. Estimates of the slowest time constant of learning given by 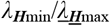 for the three scaling conditions in Experiment 1, and **g**. the two scaling conditions in Experiment 2. We generated the estimates using the Hessian matrix of networks with sigmoidal hidden units for the 15 simulated participants (eq. A6.11). Bars indicate the mean, and error bars the standard deviation. **f**. Simulated mean error in initial direction during learning in the three scaling conditions in Experiment 1 (mean of the 15 simulated participants), and **h**. in Experiment 2. The simulation results were fitted to the mean experimental curves by adjusting the learning rate *η*_I_ in eq. 6 for each simulated participant. Error bars indicate the standard error.

### High anisotropy in target distributions is associated with lower degrees of adaptation to visuomotor transformations

In experiment 2 (anisotropic target distribution task), we found that the round target layout was associated with a statistically significantly larger degree of adaptation to the visuomotor rotation than the elliptical target layout [s = 1.0, initial error: 27.9° (SD 3.0), final error: 18.9° (SD 6.0); s = 0.2, initial error: 27.0° (SD 4.4), final error: 23.3° (SD 6.3); *p* = 0.005, paired *t* test]. Figure 4d shows the time course of learning in all conditions of experiment 2. We found that the shape of the target distribution had a statistically significant effect on the learning rate parameter of each participant’s learning curve. The median of the learning rate parameter was greater for the circular target layout than for the elliptical target layout (Figure 4c) [s = 1.0, learning rate: 0.0186 (SD 0.0196); s = 0.2, learning rate: 0.0048 (SD 0.0075); *p* < 10^-3^, Wilcoxon signed rank test (data in condition s = 1.0 not normal according to Shapiro-Wilk test)]. The differences in the extent of adaptation and learning rates in all the experimental conditions suggest faster learning rates for tasks with rounder target distributions.

### Hessian analysis predicts the effects of the manipulability ellipse and target distribution on the speed of learning a visuomotor transformation

We computed the Hessian of the network defined by the computational models of 15 simulated participants to estimate the slowest time constant of learning for each condition in Experiments 1 and 2 using the distal learning and direct inverse modeling architectures. We focused on the case where the motor command ***m*** corresponds to muscle synergy activations. For distal learning, we used eq. A6.11, considering sigmoidal hidden activation functions (to use a musculoskeletal model) and a cost function that minimizes a trade-off between error and effort to compute the Hessian and its eigenvalues (Appendix 6). Results using the direct inverse modeling architecture are provided in Appendix 1. Consistent with our hypothesis, derived from the assumption of a network with linear hidden activation functions, the estimated slowest time constant of learning is larger for conditions with rounder manipulability ellipsoids or target distributions in the non-linear network (Figure 4e,g) [*λ*_***H***min_/*λ*_***H***max_ in anisotropic scaling: s = 1.0: 0.066 (SD 0.024), s = 0.5: 0.038 (SD 0.019), s = 0.2: 0.012 (SD 0.008); *λ*_***H***min_/*λ*_***H***max_ in anisotropic target distribution: s = 1.0: 0.066 (SD 0.024), s = 0.2: 0.0029 (SD 0.010)]. The measured learning rates in the experiments show a similar pattern to the estimated slowest time constant of learning 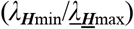 in both the distal learning and direct inverse modeling architectures, with a monotonic relationship between the scaling factor of the manipulability ellipsoid and target distributions, and the observed speed of learning (Figure 4a,c).

Furthermore, for distal learning, the simulated learning curves in both the anisotropic scaling and target distribution tasks (Figure 4f,h) closely match the learning curves observed experimentally after selecting an appropriate learning rate η_I_ (eq. 6) for each simulated subject (anisotropic scaling task: R^2^ = 0.90, anisotropic target distribution task: R^2^ = 0.90). Note, however, that the simulated learning curves using direct inverse modeling (Figure A1.2c) have a lower quality fit to the experimental observations (anisotropic scaling: *R*^2^ = 0.78, anisotropic target distribution: *R*^2^ = 0.83).

### Hessian analysis predicts differences in learning rates in virtual surgery tasks

We applied the computational framework to a virtual surgery task [23]. We found that the distal learning framework can only predict differences in the learning rates of an incompatible and a compatible virtual surgery when the model includes explicitly defined muscle synergies [distal learning, non-linear network; *λ*_***H***min_/*λ*_***H***max_ with synergies, compatible: 0.029 (SD 0.016); incompatible: 0.017 (SD 0.014); *λ*_***H***min/_*λ*_***H***max_ of compatible larger for 13 of the simulated participants] (Figure 5a). If muscle synergies are not included in the model, the framework predicts that the learning speed in the incompatible task is slightly larger than in the compatible task, which does not align with experimental observations [distal learning, non-linear network; *λ*_***H***min_/*λ*_***H***max_ without synergies, compatible: 0.094 (SD 0.036); incompatible: 0.112 (SD 0.048); *λ*_***H***min_/*λ*_***H***max_ of incompatible larger for 12 of the simulated participants]. In fact, for linear networks without muscle synergies, there is no difference in the maximum speed of learning between the compatible and incompatible tasks. This is because the eigenvalues of ***P***^T^***P*** are not a function of ***T***, given that for non-zero eigenvalues *λ*(***P***^T^***P***) = *λ*(***PP***^T^), and ***PP***^T^ = ***MTT***^T^***M***^T^ = ***MM***^T^, as by definition, **T** is an orthonormal matrix. However, the direct inverse modeling framework predicts differences in the learning rates of both tasks whether muscle synergies are explicitly defined in the model or not [direct inverse modeling, *λ*_***H***min_/*λ*_***H***max_ without synergies, compatible: 0.05 (SD 0.039); incompatible: 0.007 (SD 0.006); *λ*_***H***min_/*λ*_***H***max_ of compatible larger for 14 of the simulated participants] (Figure A1.2a). In both simulated frameworks, the simulated learning curves in the compatible and incompatible surgeries closely match the learning curves observed experimentally after selecting an appropriate learning rate η_I_ for each simulated subject (distal learning: *R*^2^ = 0.88; direct inverse modeling: *R*^2^ = 0.86) (Figure 5b and A1.2c).

**Figure 5.**
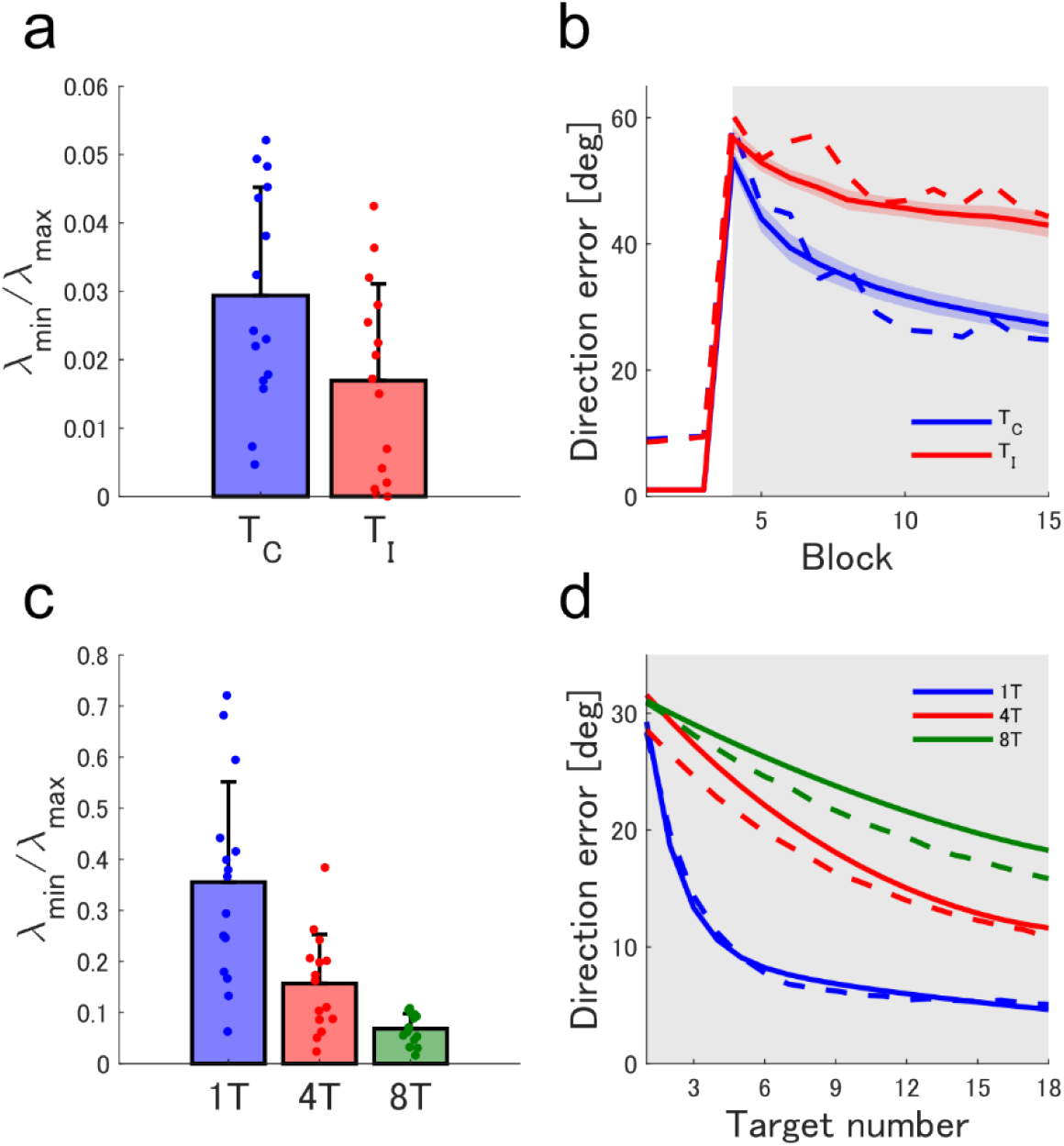
Simulation results in the virtual surgery and multi-target tasks. **a**. Estimates of the slowest time constant of learning given by *λ*_***H***min_/*λ*_***H***max_ for the compatible (***T***_C_) and incompatible (***T***_I_) virtual surgeries. We generated the estimates using networks with sigmoidal hidden units and muscle synergies in the distal learning framework for the 15 simulated participants (eq. A6.11). Bars indicate the mean, and error bars the standard deviation. **b**. Simulated mean error in initial direction during learning in ***T***_C_ and ***T***_I_. Simulated learning curves were fitted to the mean experimental curves by adjusting the learning rate *η*_I_ in eq. 6. Solid lines correspond to the mean error in the 15 simulated participants. Dashed lines correspond to the mean error in experimental observations [23]. Error bars indicate the standard error. **c**. Estimates of the slowest time constant of learning given by *λ*_***H***min_/*λ*_***H***max_ for the multi target task with 1, 4 and 8 targets (1T, 4T and 8T). We generated the estimates as indicated in **a**. **d**. Simulated mean error in initial direction during learning in 1T, 4T and 8T. Learning curves are double exponentials fitted to the mean simulated curves, which in turn were fitted to the mean experimental curves by adjusting the learning rate *η*_I_ in eq. 6. Dashed lines correspond to double exponential fits to the experimental observations [30].

### Hessian analysis predicts differences in learning rates in multi-target task

Finally, we applied the computational framework to a visuomotor rotation task in which experimental conditions are defined by the number of targets in the task (2, 4 or 8 targets) [30]. We found that the framework predicts that the speed of learning decreases as the number of targets increases, consistent with experimental observations [distal learning, non-linear network; *λ*_***H***min_/*λ*_***H***max_ with synergies, 1 target: 0.356 (SD 0.196); 4 targets: 0.157 (SD 0.096); 8 targets: 0.069 (SD 0.029); *λ*_***H***min_/*λ*_***H***max 2T_ > *λ*_***H***min_/*λ*_***H***max 4T_ and *λ*_***H***min_/*λ*_***H***max 4T_ > *λ*_***H***min_/*λ*_***H***max 8T_ for all 15 simulated participants] (Figure 5c). The direct inverse modeling framework also predicts a monotonic relationship between the number of targets and the speed of learning. However, the monotonic pattern of this relationship is appreciably different from the experimental observations (Figure A1.2a). The simulated learning curves in the number of targets task with the distal learning architecture closely match the learning curves observed experimentally after selecting an appropriate learning rate *η*_I_ for each simulated subject (*R*^2^ = 0.92) (Figure 5d). However, the simulated curves using direct inverse learning have a lower quality fit to the experimental observations (*R*^2^ = 0.77) (Figure A1.2c).

## Discussion

In this study, we showed that analyzing models of human motor learning through the lens of optimization theory allows us to predict how fast the CNS can learn new motor tasks. Specifically, we analyzed learning in a set of isometric force tasks for two different inverse learning paradigms, distal learning and direct inverse modeling. The simplicity of the forward physics of the isometric tasks allowed us to analytically derive the Hessian of a linear approximation of the learning system. We found that the eigenvalues of the Hessian matrix, and thus the speed of learning, are a function of the shapes of the force manipulability ellipsoid of the arm and the distribution of target forces. We extended these theoretical results to a non-linear representation of the motor system that includes muscles. Furthermore, our modeling framework establishes that these results are generalizable to other task conditions, and even different isometric tasks such as virtual surgeries. Although in the main text we have used backpropagation to illustrate the properties of the speed of learning under gradient-following algorithms, we also showed that our results do not depend on this choice and are also valid for the more biologically plausible node perturbation algorithm (Appendix 2).

### Testable predictions

Based on our theoretical results, we hypothesized that altering the manipulability ellipsoid of the arm or the force target distributions would directly impact the learning speed during a visuomotor rotation (VMR) in the isometric reaching task. We tested our hypothesis by having participants perform the VMR task under four different conditions. In the first three conditions, we altered the roundness of the manipulability ellipsoid in a virtual environment. In the fourth condition we used a target distribution in the shape of an ellipse. We found that rounder manipulability ellipsoids and target distributions allow for faster learning in the VMR task, confirming our hypothesis.

Notice, however, that we have made a simplifying assumption regarding the role of the target distribution. The eigenvalues of the Hessian matrix of the distal learning system are a function of the distribution of RBF inputs to the network rather than simply the two-dimensional distribution of target forces (eq. 17). Therefore, variables such as the number, centers and shapes of the RBF units also impact the eigenvalues of the Hessian. Unfortunately, these properties of the RBFs are hard or impossible to control in an experimental setup. However, because the RBF units encode the desired forces, the distribution of RBF inputs must also encode information about the distribution of target forces. This makes the distribution of target forces an ideal proxy for the distribution of RBF inputs, as it is an easily controllable variable.

Our results suggest that the differences in the speed of learning in the task brought about by factors such as manipulability ellipsoids and target distributions may be an inherent computational property of learning in the isometric task. Both of these factors predict differences in the speed of learning for both the distal learning and direct inverse modeling paradigms, indicating that the differences may arise regardless of a particular architecture or optimized cost function (performance error vs motor error). However, although both inverse learning models are affected in a qualitatively similar way by these factors, the predicted relative speed of learning in different tasks is quantitatively different (Figure 4eg, Figure 5ac, A1. 1a), which is reflected in the simulated learning curves. Other factors seem to be particular to a given architecture. For example, the activation levels of muscles are involved in shaping the Hessian and thus the speed of learning in distal learning (eq. A6.4), but not in direct inverse modeling. Similarly, another architecture-specific factor that could influence the speed of learning is the nature of the RBFs encoding the network inputs, as mentioned above. Therefore, our framework can be used as a tool to generate systematic hypotheses to infer the type of algorithms that the CNS uses for motor learning in different tasks. For instance, our results show that the distal learning paradigm is able to describe the relative speeds of learning in all of the VMR tasks more adequately with respect to the data than the direct inverse learning paradigm. However, both paradigms provide similarly adequate descriptions of the speed of learning in the virtual surgery task.

Some of the factors that influence the speed of learning identified by our framework may be obvious based on previous theoretical and experimental studies. For example, the number of targets in a VMR task has been shown to affect the speed of learning because learning of rotations is only generalized locally. Learning around one target does not contribute to learning around faraway targets, slowing down learning overall [30]. Since the models we used encode the positions of targets with local RBFs, it is not surprising that our simulations can reproduce the effects of the number of targets on the speed of learning. However, it is notable that this effect is observable by examining the Hessian of the learning models. Increasing the number of targets also increases the rank of the correlation matrix of the RBF inputs, resulting in a decrease in the ratio of the smallest and largest eigenvalues of the matrix, which is associated with slower learning. This suggests that other unexplored factors that affect the speed of learning could be identified by analyzing the Hessian of the learning models, even if an intuitive reason for the effect is elusive.

Additionally, our framework could potentially allow us to generate hypotheses about the role of different structures in the CNS on the speed of learning. Here, we found that a distal learning approach requires muscle synergies to be actual neural structures in the CNS to reproduce the results of virtual surgery experiments (Appendix 7). These neural circuits in conjunction with the virtual surgeries reshape the manipulability ellipsoid of the arm in the virtual environment, and thus, compatible and incompatible surgeries elicit a difference in learning speeds. Otherwise, without these neural structures, the virtual surgeries have an unspecified effect on the manipulability ellipsoid, and the differences between compatible and incompatible surgeries are not captured by the model (see in-line equations in *Hessian analysis predicts differences in learning rates in virtual surgery tasks*). In contrast, the direct inverse modeling approach can reproduce the experimental results with or without neural structures for muscle synergies in the model. This is because the virtual surgeries themselves differentially change the distribution of realized forces, which influences the speed of learning (Appendix 1).

Note, however, that care must be taken when comparing the results on the speed of learning in different architectures. Although both inverse learning paradigms, distal learning and direct inverse modeling, rely on gradient descent to learn an inverse, the way in which the inverse model is updated in both paradigms is different. Distal learning attempts to minimize predicted errors on performance via a forward model of the physics of the body, whereas direct inverse modeling attempts to minimize the error between the realized motor command and the motor command it would use to produce the realized action. That is, the quantities optimized in both paradigms are different. Therefore, our analysis pertains only to the learning rates of the quantity optimized in each paradigm. Our results are directly applicable in the case of distal learning, as the performance error is one of the main variables of interest in motor learning. In contrast, in the case of direct inverse modeling, our analysis corresponds to the learning rate in the motor error, which in general is not possible to measure experimentally, and does not directly relate to task performance. However, previous studies have shown that in linear systems, minimization of the motor error also leads to minimization of the performance error given appropriate initial conditions and/or exploration noise [43]. Furthermore, our direct inverse modeling simulations show that learning rates in the motor and performance domains correlate across tasks (Figure A1. 2), so our analysis is still useful provided that the minimization of performance and motor errors is coupled. This condition would be violated in the case of failure to form an adequate inverse model [11] or the occurrence of other failure modes [9].

### Future perspectives

One open question is the relation between the timescales of learning given by the eigenvalues of the Hessian and the different timescales of learning processes identified in the CNS. Motor adaptation is often described as the superposition of slow and fast processes that allow fast error corrections in the face of a changing environment and retention of the learned task in a stable environment [44]. Models employing one fast and one slow process have been quite successful in describing motor adaptation in many tasks [45,46], although processes with multiple timescales have been identified [47]. The neural basis of these processes is still not well understood, but may originate from different circuits in the CNS [47,48]. However, in our framework, the multiple timescales defined by the Hessian describe the inherent dynamics of learning as an optimization process, without separate neural circuits operating at different timescales, as discussed below. Therefore, further research that considers the inherent dynamics of learning to study learning processes with different timescales seems necessary.

The dynamics of learning an inverse model are characterized by the eigenvalues of the Hessian and are reflected in the resulting learning curves as different timescales. The number of non-zero eigenvalues of the Hessian (its rank) then defines the number of timescales involved in learning. In the distal learning model, the eigenvalues of the Hessian are defined by the eigenvalues of the ***P***^T^***P*** and ***ϕϕ***^T^ matrices, representing the manipulability ellipsoid and target distribution, respectively (eq. 16). ***P***^T^***P*** is in general only rank 2, whereas the rank of ***ϕϕ***^T^ can be as large as the number of RBF input units, and depends on the number of targets. In our simulations, the rank of the Hessian is in general 16, as most experiments modeled use 8 targets (rank(***H***) = rank(***P***^T^***P***)·rank(***ϕϕ***^T^)). We found that considering only the ratio of the slowest and fastest time constants predicts the relative speed of learning of several tasks quite well. However, it is possible that an analysis of other timescales (i.e., other eigenvalues) could provide a more specific description of the learning curves in the tasks. This could be especially important in tasks in which the time frame of the experiment does not allow convergence to zero error, as is the case in some of the experiments we conducted here. Additionally, considering other timescales could provide further differentiation between candidate models to describe learning in a task.

### Limitations

One limitation of our framework is that it assumes that updates to the inverse model are obtained based on the batch gradient, which attempts to minimize the error for all target forces simultaneously. That is, errors for the whole set of targets are known and used for each learning update. However, in humans, learning takes place mainly on a trial-by-trial basis, although a memory of previous errors can contribute to learning [49]. This shorter learning timeframes are more akin to online or stochastic gradient descent. Therefore, because the batch and the singletrial gradient are different in general, the actual limit of the speed of learning in the model may differ from the theoretical estimates. In practice, however, we found that using this assumption still adequately predicts the relative learning speed in different tasks both in simulations that use online learning, and experiments.

Another limitation comes from our choices in the experimental design. Due to restrictions arising from the COVID-19 pandemic, we conducted the experiments using an online platform [37]. Although the results obtained from the online platform are similar to in-person results for VMR tasks [37–39], the arm posture in our experiment should be controlled, as it is related to the shape of the manipulability ellipse of the arm because the posture determines the arm Jacobian [25]. We instructed the participants to adopt a similar posture across all experimental conditions, but did not control for the posture. However, considering that changes in arm posture during the experiment are unlikely to have as dramatic an effect on the manipulability ellipse as the visuomotor transformations that we used, any effects of changes in arm posture are likely not important.

An additional limitation stemming from our choices on experimental design is that our computational framework considers an isometric task in which the forward physics of the arm are linear. We referred to the experimental task as “quasi-isometric”, as the cursor movements are controlled by small finger and/or arm movements (ideally around 1 cm in end-point movement extent, without appreciable changes in arm joint angles). Therefore, even though the forward physics of the arm are not linear because of these movements, the movement extent is small enough that a local linear approximation may be adequate for its description. Nevertheless, previous research in motor control and learning in arm reaching tasks shows that modeling reaching tasks as a linear system can have good descriptive and predictive power [44].

### Conclusion

In conclusion, we have developed a framework to theoretically identify factors that influence the speed of learning. This could potentially be used in more complex tasks to systematically identify body and environment parameters that facilitate learning, as well as to evaluate the difficulty of the task based on the elements of the task. Additionally, it could be used to systematically find candidate hypotheses about model architecture and components in the CNS to describe learning in a task. Our results show the relevance of the theory of artificial neural networks to understand the mechanisms underlying learning in the CNS, as advocated in recent perspectives [50–52].

## Supporting information

Appendix 1. Direct inverse modeling

Appendix 2. Node perturbation

Appendix 3. The shape of the cost function and the speed of learning

Appendix 4. Gradient and Hessian matrix in a linear network

Appendix 5. Hessian matrix in a network that minimizes error and effort

Appendix 6. Hessian matrix in a network with non-linear hidden units

Appendix 7. Muscle synergies in simulation

Appendix 8. Creation of simulated participants

Appendix 9. Table of variables

## Notes

### Competing Interest Statement

The authors have declared no competing interest.

